# In and outs of Chuviridae endogenous viral elements: origin of a retrovirus and signature of ancient and ongoing arms race in mosquito genomes

**DOI:** 10.1101/2020.02.15.950899

**Authors:** Filipe Zimmer Dezordi, Crhisllane Rafaele dos Santos Vasconcelos, Antonio Mauro Rezende, Gabriel Luz Wallau

## Abstract

**Background:** Endogenous viral elements (EVEs) are sequences of viral origin integrated into the host genome. EVEs have been characterized in various insect genomes, including mosquitoes. A large EVE content has been found in *Aedes aegypti* and *Aedes albopictus* genomes among which a recently described *Chuviridae* viral family is of particular interest, owing to the abundance of EVEs derived from it, the discrepancy in the endogenized gene regions and the frequent association with retrotransposons from the BEL-Pao superfamily. In order to better understand the endogenization process of chuviruses and the association between chuvirus glycoproteins and BEL-Pao retrotransposons, we performed a comparative genomics and evolutionary analysis of chuvirus-derived EVEs found in 37 mosquito genomes.

**Results:** We identified 428 EVEs belonging to the *Chuviridae* family confirming the wide discrepancy between the number of genomic regions endogenized: 409 glycoproteins, 18 RNA-dependent RNA polymerases and one nucleoprotein region. Most of the glycoproteins (263 out of 409) are associated specifically with retroelements from the Pao family. Focusing only on well assembled Pao retroelement copies, we estimated that 263 out of 379 Pao elements are associated with chuvirus-derived glycoproteins. Seventy-three potentially active Pao copies were found to contain glycoproteins into their LTR boundaries. Thirteen out of these were classified as complete and likely autonomous copies, with a full LTR structure and protein domains. We also found 116 Pao copies with no trace of glycoproteins and 37 solo glycoproteins. All potential autonomous Pao copies, contained highly similar LTRs, suggesting a recent/current activity of these elements in the mosquito genomes.

**Conclusion:** Evolutionary analysis revealed that most of the glycoproteins found are likely derived from a single or few glycoprotein endogenization events associated with a recombination event with a Pao ancestral element. A potential fully functional Pao*-*chuvirus hybrid (named Anakin) emerged and the glycoprotein was further replicated through retrotransposition. However, a number of solo glycoproteins, not associated with Pao elements, can still be found in some mosquito genomes 114 million years later, suggesting that these glycoproteins were likely domesticated by the host genome and may participate in an antiviral defense mechanism against both chuvirus and Anakin retrovirus.

## Background

Viruses have long-term and intricate interactions parasitizing host cells and hence both viruses and host are subject to an endless arms race (FORTERRE; PRANGISHVILI, 2009). A large body of evidence currently supports that virus/host interactions can occur at both the protein and nucleic acid levels. One clear example of the last, is that viral genomic sequences can be integrated into the host genome (FESCHOTTE; GILBERT, 2010; JOHNSON, 2019). The process of viral genome integration is called endogenization and normally occurs as a “life cycle” stage in viral groups such as retroviruses and phage DNA viruses (WEISS, 2016). However, recent studies have shown that genomes, or genomic regions, of non-integrative viruses can also be found integrated into various eukaryotic genomes (FESCHOTTE; GILBERT, 2010; JOHNSON, 2019; KATZOURAKIS; GIFFORD, 2010). These viral loci have been called endogenous viral elements (EVEs).

Endogenization can occur by two main mechanisms: through non-homologous recombination mediated by double-strand break repair pathway of the host cell; or mediated by proteins, such as reverse transcriptases and integrases, from the endogenous retrotransposons—envelope-free retrovirus-like elements (KATZOURAKIS; GIFFORD, 2010). Recent findings on mosquito genomes suggest that the latter mechanism is likely to be the most important, since the abundance and diversity of EVEs are positively correlated with retrotransposon abundance and activity (PALATINI *et al.*, 2017; WHITFIELD *et al.*, 2017). Likewise, EVEs and retrotransposon loci normally exist in close proximity in the host chromosome, suggesting a causal role between the activity of certain retrotransposon families and EVE integration (PALATINI *et al.*, 2017; WHITFIELD *et al.*, 2017). EVEs are normally found as fragments of exogenous viral genomes and it is therefore unlikely that they are able to generate new virus particles or infect new cells. Therefore, there are three main hypotheses regarding the fate and impact of these elements on the host genome: i) EVEs may evolve neutrally accumulating mutations and degenerate over time; ii) EVEs may be co-opted by the host genomes, giving rise to new functional host genes; and iii) EVEs may play an antiviral role, generating small RNAs which degrade cognate exogenous viral RNA or non-functional viral proteins that hinders proper assembly/maturation of a new viral particle or blocks the viral receptor on the host cell surface (ARMEZZANI *et al.*, 2014; ITO *et al.*, 2013; ROBINSON *et al.*, 1981). The vast majority of studies on EVEs found in mosquito genomes focus on the role of piRNA production as a post-transcriptional regulatory mechanism of exogenous circulating viruses (PALATINI *et al.*, 2017; WHITFIELD *et al.*, 2017; THÉRON *et al.*, 2013). On the other hand, there are few studies of the role of EVE proteins in the antiviral response or their role in the emergence of new host genes.

*Chuviridae* is a recently discovered RNA viral family of negative-sense single-stranded viruses characterized by metatranscriptomic and bioinformatic analysis only (SHI *et al.*, 2016). The information available on this family is limited to its distribution (it likely infects several arthropod species including mosquitoes), its variable genomic structure (unsegmented, bisegment and circular), and the presence of a number of EVEs in species from Amphipoda, Hemiptera, Coleoptera, Hymenoptera and Diptera genomes (Shi et al. 2016). An in-depth descriptive analysis of chuvirus-derived EVEs exists only for the *Ae. aegypti* mosquito genome (WHITFIELD *et al.*, 2017), in which they displayed three intriguing features: a great abundance of chuvirus-derived EVEs compared to other viral families (it is the second most abundant EVE family, outnumbered only by the *Rhabdoviridae* family); an association with retroelements from the BEL-Pao superfamily; and a striking difference in the viral genome fragment endogenized - a much higher quantity of glycoprotein (Gly) compared to RNA dependent RNA polymerase (RdRp) and nucleoprotein (NP) sequences (WHITFIELD *et al.*, 2017). Endogenization of different viral regions are expected to be influenced by the virus genome structure and orientation of the RNA replication process. Whitfield et al., 2017 have shown that EVEs derived from another non-segmented negative-sense single-stranded virus from the *Rhabdoviridae* family occur in the following order of abundance: NP -> Gly -> RdRp ressambling the order in which the viral genomes is replicated in this viral family. On the other hand, *Chuviridae* viruses, which are similar to *Rhabdoviridae* in genomic structure (NP-Gly-RdRp), shows a very different endogenization pattern, with eighty-seven endogenous Gly sequences, only four from RdRp and no endogenized nucleoproteins (WHITFIELD *et al.*, 2017). It may indicate that chuvirus-derived EVEs found in *Ae. aegypti* genome were either derived from a exogenous chuvirus with segmented or non-segmented genome and for some reason Gly has been majorly endogenized, or that the endogenization of chuvirus genomic regions occurred evenly at the beginning but only Gly was maintained through the evolutionary time.

Endogenous repetitive elements are abundant in metazoan genomes and mosquito genomes from the *Aedes* genus are particularly full of retrotransposons/retroviruses (GOUBERT *et al.*, 2015; NENE *et al.*, 2006; WHITFIELD *et al.*, 2017). One of the most abundant LTR retrotransposons/retroviruses are from the BEL-Pao superfamily which is also very abundant and widespread in other metazoan genomes (MALIK, H. S.; HENIKOFF; EICKBUSH, 2000). Elements from this group have two long terminal repeat (LTR) regions, consisting of between 100 and 900 base pairs, and two coding regions — a capsid protein with a GAG domain, and a polyprotein, which commonly has four domains: aspartic protease (PR), reverse transcriptase (RT), RNAse H (RH) and integrase (INT). Moreover, at least three elements (Roo, Tas and Cer-7) have an envelope-like protein (Gly) downstream of the polyprotein, which were acquired from *Gypsy*, Phlebovirus and Herpesvirus, respectively (FELDER *et al.*, 1994; BROWNING *et al.*, 1996; MALIK, HARMIT S.; HENIKOFF, 2005).

In view of all the intriguing aforementioned chuvirus/BEL-Pao/host features, we investigated in depth which biological phenomenon has generated the high abundance of chuvirus glycoproteins found in mosquito genomes and examined the role these glycoproteins may play in BEL-Pao retroelements and mosquito biology. Our results showed, for the first time, that these EVEs are broadly present in mosquito genomes and that a large majority of the glycoproteins are physically associated with elements from the Pao family. We also found that most of the chuvirus-derived glycoproteins are structurally associated with potentially autonomous Pao elements and are likely to play a role in viral particle formation as an envelope protein. However, we also found structurally conserved solo glycoproteins, suggesting a potential role for these in antiviral defense mechanisms via receptor competition against *bona fide* chuviruses and/or the new retrovirus (named Anakin in this paper) emerging from the acquisition of the chuvirus glycoprotein by Pao retroelements.

## Methods

### Data Collection

A non-redundant database of proteins including all taxa and a database of chuvirus genomes were obtained from NCBI (last accessed January 2018). Mosquito genomes were retrieved from NCBI and Vectorbase (last accessed January 2018) and chuvirus-derived EVEs already identified in mosquito genomes were retrieved from (WHITFIELD *et al.*, 2017). Mosquito genome sources and assembly metrics are presented in **Supplementary Material 1**. All command lines used in the present study can be found in **Supplementary Material 2**.

### EVE Screening

Two BLAST-based strategies were used. The first used the chuvirus genome dataset as a query in a tBLASTx (ALTSCHUL *et al.*, 1989) screening against mosquito genomes, and the second used the EVEs from (WHITFIELD *et al.*, 2017) as a query in a BLASTn analysis against mosquito genomes. All EVE regions were extracted from mosquito genomes in two ways: I - only the ungapped aligned region; II - ungapped aligned region plus 10kb of each flanking region.

The resulting sequences were used as a query in a BLASTx analysis against the non-redundant protein database with different filters in order to eliminate false positives and false negatives in chuvirus EVE identification. This was done for two reasons. First, as a number of viral proteins are still not annotated in the databases and hence if one considers the first hit alone in order to determine the viral origin one would rule out some EVEs and produce a false negative result. Second, some wrongly annotated viral proteins would cause the first hit to be mis-annotated as viral, while the following two or more hits would indicate that the sequence belongs to another taxon, generating a false positive result. Queries that showed four or fewer matches were annotated as an EVE if the best match was a viral protein. For queries with five or more subject matches, the proportion of the five best matches was taken into consideration, in accordance with the following criterion: a sequence is annotated as an EVE only if three or more subjects have viral proteins.

The EVE sequences containing the flanking sequences were clusterized using CD-HIT (LI; GODZIK, 2005) to remove redundancy. Flanking sequences were retained to avoid clusterization of identical or very similar copies while allowing the removal of the same EVE copy recovered using the two search strategies (chuvirus genomes and EVEs).

### Chuvirus EVE characterization

Using the above strategy, we obtained a total of 428 chuvirus-derived EVEs from mosquito genomes, of which 409 were glycoproteins, 18 RNA-dependent RNA polymerases and one nucleoprotein region. However, a number of these were found in small-size contigs and in close proximity to indeterminate “NNN” regions in the assembly. In order to avoid genome assembly problems, we restricted our analysis to contigs bearing EVEs with at least 4 kb (LLORENS *et al.*, 2011) of each flanking region, with no undetermined bases in these regions (**Supplementary Material 3**).

For the remaining 322 sequences, we extracted potential coding regions using the EMBOSS *getorf* (http://www.bioinformatics.nl/cgi-bin/emboss/getorf) tool, and ORFs containing fewer than 100 amino acids were removed from further steps, resulting in 279 sequences.

### Flanking sequence analysis

Once they had been passed through the aforementioned filters, all EVEs plus flanking regions were translated using the EMBOSS *getorf* software, and the resulting ORFs were used in a domain-signature analysis with BATCH-CDD (MARCHLER-BAUER *et al.*, 2011) to characterize the genomic context of each EVE loci. Sequences with BEL-Pao signature domains — GAG, PR, RT, RNAse H, and INT (whether or not flanked by long terminal repeats - LTRs) — were considered to be putative hybrids of a BEL-Pao element and chuvirus-derived sequences. For graphical representation, genetic maps are constructed with karyoploteR R-package (GEL; SERRA, 2017).

### Search for all homologous BEL-Pao elements

Nucleotide sequences of complete BEL-Pao elements containing domain signatures and LTRs recovered in the previous analysis were used as queries in a BLASTn analysis against the respective mosquito genomes to recover all homologous BEL-Pao copies. Sequences retrieved in this step were recovered with 10kb of each flanking region and screened for the presence of chuvirus-derived EVEs. Long terminal repeat (LTR) regions from all sequences retrieved were evaluated using LTR_FINDER and default parameters (XU; WANG, 2007).

All EVEs were sorted, based on RdRp, Gly or NP proteins and BEL-Pao Retrotransposons based on whether they are associated with chuvirus-derived glycoproteins or not and the structural conservation of the copies, into the following categories: i) potentially active retrotransposons containing both LTRs, BEL-Pao conserved protein domains (GAG-PR-RT-RH-INT) with or without a chuvirus-derived glycoprotein; ii) defective elements which have at least one BEL-Pao domains and one or no LTRs; iii) solo LTRs; and iv) solo glycoproteins (**Table 1**).

**Table 1:** General features of chuvirus-like proteins and BEL-Pao retroelements identified.

| Genome | Chuvirus |  |  |  | With Glyco* |  |  |  | Without Glyco* |  |  |  | Solo-LTR |
| --- | --- | --- | --- | --- | --- | --- | --- | --- | --- | --- | --- | --- | --- |
|  | Nucleoprotein | Solo-Glyco** | Glyco+Pao | RdRp | Complete | Defective+LTR | Defective | Total | Complete | Defective+LTR | Defective | Total |  |
| *Aedes aegypti* AaegL3 | 0 | 5/4 | 23 | 2 | 1 | 2 | 20 | 23 | 0 | 9 | 15 | 24 | 9 |
| *Aedes aegypti* Aag2 | 1 | 15/8 | 55 | 4 | 0 | 20 | 35 | 55 | 0 | 9 | 9 | 18 | 14 |
| *Aedes aegypti* BV | 0 | 14/0 | 1 | 4 | 0 | 0 | 1 | 1 | 0 | 0 | 1 | 1 | 0 |
| *Aedes albopictus* Rimini | 0 | 20/0 | 1 | 2 | 0 | 0 | 1 | 1 | 0 | 0 | 0 | 0 | 0 |
| *Aedes albopictus* Foshan | 0 | 19/3 | 27 | 3 | 0 | 2 | 25 | 27 | 0 | 1 | 18 | 19 | 4 |
| *Aedes albopictus* C636 | 0 | 3/3 | 85 | 3 | 9 | 15 | 61 | 85 | 2 | 20 | 28 | 50 | 82 |
| *Culex quinquefasciatus* | 0 | 3/3 | 17 | 0 | 3 | 9 | 5 | 17 | 0 | 0 | 0 | 0 | 20 |
| *Anopheles albimanus* | 0 | 0 | 3 | 0 | 0 | 0 | 3 | 3 | 0 | 0 | 0 | 0 | 0 |
| *Anopheles arabiensis* | 0 | 2/0 | 2 | 0 | 0 | 0 | 2 | 2 | 0 | 0 | 0 | 0 | 0 |
| *Anopheles atroparvus* | 0 | 4/0 | 1 | 0 | 0 | 0 | 1 | 1 | 0 | 0 | 0 | 0 | 0 |
| *Anopheles christyi* | 0 | 0 | 0 | 0 | 0 | 0 | 0 | 0 | 0 | 0 | 0 | 0 | 0 |
| *Anopheles coluzzi* | 0 | 1/0 | 1 | 0 | 0 | 0 | 1 | 1 | 0 | 0 | 0 | 0 | 0 |
| *Anopheles culicifacies* | 0 | 0 | 0 | 0 | 0 | 0 | 0 | 0 | 0 | 0 | 0 | 0 | 0 |
| *Anopheles darlingi* | 0 | 0 | 0 | 0 | 0 | 0 | 0 | 0 | 0 | 0 | 0 | 0 | 0 |
| *Anopheles dirus* | 0 | 5/2 | 5 | 0 | 0 | 0 | 5 | 5 | 0 | 0 | 0 | 0 | 0 |
| *Anopheles epiroticus* | 0 | 4/0 | 3 | 0 | 0 | 0 | 3 | 3 | 0 | 0 | 0 | 0 | 0 |
| *Anopheles farauti* | 0 | 5/0 | 2 | 0 | 0 | 0 | 2 | 2 | 0 | 0 | 0 | 0 | 0 |
| *Anopheles funestus* | 0 | 7/1 | 1 | 0 | 0 | 0 | 1 | 1 | 0 | 0 | 0 | 0 | 0 |
| *Anopheles gambiae* PEST | 0 | 0 | 2 | 0 | 0 | 0 | 2 | 2 | 0 | 0 | 0 | 0 | 0 |
| *Anopheles gambiae* Pimpirena | 0 | 0 | 6 | 0 | 0 | 1 | 5 | 6 | 0 | 0 | 0 | 0 | 1 |
| *Anopheles gambiae* S | 0 | 3/3 | 1 | 0 | 0 | 0 | 1 | 1 | 0 | 0 | 0 | 0 | 0 |
| *Anopheles gambiae* BV | 0 | 0 | 0 | 0 | 0 | 0 | 0 | 0 | 0 | 0 | 0 | 0 | 0 |
| *Anopheles koliensis* | 0 | 1/0 | 1 | 0 | 0 | 0 | 1 | 1 | 0 | 0 | 0 | 0 | 0 |
| *Anopheles maculatus* m3 | 0 | 3/0 | 0 | 0 | 0 | 0 | 0 | 0 | 0 | 0 | 0 | 0 | 0 |
| *Anopheles maculatus* BtQ1 | 0 | 12/3 | 15 | 0 | 0 | 1 | 14 | 15 | 0 | 0 | 0 | 0 | 4 |
| *Anopheles melas* | 0 | 1/0 | 0 | 0 | 0 | 0 | 0 | 0 | 0 | 0 | 0 | 0 | 0 |
| *Anopheles merus* | 0 | 1/1 | 1 | 0 | 0 | 0 | 1 | 1 | 0 | 0 | 1 | 1 | 0 |
| *Anopheles minimus* | 0 | 4/2 | 1 | 0 | 0 | 0 | 1 | 1 | 0 | 0 | 0 | 0 | 0 |
| *Anopheles nili* | 0 | 1/0 | 2 | 0 | 0 | 0 | 2 | 2 | 0 | 0 | 0 | 0 | 0 |
| *Anopheles punctulatus* | 0 | 0 | 0 | 0 | 0 | 0 | 0 | 0 | 0 | 0 | 0 | 0 | 0 |
| *Anopheles quadriannulatus* | 0 | 3/0 | 2 | 0 | 0 | 0 | 2 | 2 | 0 | 0 | 0 | 0 | 0 |
| *Anopheles sinensis* SINENSIS | 0 | 1/1 | 1 | 0 | 0 | 0 | 1 | 1 | 0 | 0 | 0 | 0 | 0 |
| *Anopheles sinensis* china | 0 | 5/1 | 0 | 0 | 0 | 0 | 0 | 0 | 0 | 0 | 0 | 0 | 0 |
| *Anopheles stephensi* Indian | 0 | 1/1 | 1 | 0 | 0 | 0 | 1 | 1 | 0 | 0 | 2 | 2 | 0 |
| *Anopheles stephensi* SDA | 0 | 1/1 | 2 | 0 | 0 | 0 | 2 | 2 | 0 | 0 | 1 | 1 | 0 |
| *Anopheles cracens* | 0 | 0 | 0 | 0 | 0 | 0 | 0 | 0 | 0 | 0 | 0 | 0 | 0 |
| *Anopheles aquasalis* | 0 | 2/0 | 1 | 0 | 0 | 0 | 1 | 1 | 0 | 0 | 0 | 0 | 0 |
| Total | 1 | 146/37 | 263 | 18 | 13 | 50 | 200 | 263 | 2 | 39 | 75 | 116 | 134 |

\*Pao retroelements; \*\*Non-filtred/filtred. Defect means elements that do not have all domains of a complete BEL-Pao retroelement (Both LTRs, GAG protein and polyprotein with all domains).

### Molecular modeling

Molecular modeling was performed for three groups of glycoproteins: I - Solo glycoproteins, corresponding to glycoproteins without the BEL-Pao signature in their flanking regions; II - glycoproteins found inside of BEL-Pao element boundaries (LTRs) - in order to select some Gly proteins found inside BEL-Pao retrotransposons (group II), an amino acid distance matrix was created using UGENE (OKONECHNIKOV; GOLOSOVA; FURSOV, 2012). One random sequence was chosen for molecular modeling from each cluster exhibiting more than 90% similarity (**Supplementary Material 4**).; and III - glycoproteins from *bona fide* chuviruses previously characterized by metatranscriptomics in mosquitoes.

The Phyre2 server (KELLEY *et al.*, 2015) was used to select a template for each protein of interest. The 3D structure of each template selected was obtained from the Protein Data Bank (PDB) (Berman et al. 2000) and submitted, together with the Gly sequence, to the Modeller package version 9.23 multimer modeling algorithm. The predicted models were evaluated for their stereochemical parameters using the Procheck tool (Laskowski et al. 1993).

### Solo LTRs

LTRs from previously characterized elements were used as a query against mosquito genomes in a BLASTn analysis. Matches were recovered with 10 kb of flanking region. Sequences with a single LTR matching the mean length of LTRs identified using full BEL-Pao elements (around 670 base pairs) and no surrounding BEL-Pao domains were considered to be Solo LTRs (**Supplementary Material 5**).

### Evolutionary analysis

Two analyses were performed. The first was based on Gly protein sequences using only amino acid sequences with more than 100 residues. This analysis included most EVEs identified and chuvirus Gly from the literature and from the NCBI. The second analysis involved the evolutionary history of BEL-Pao retrotransposons and used amino acid sequences from reverse transcriptase and RNAse H (the most abundant domains present on BEL-Pao as characterized in the present study) (**Supplementary Material 6**). This second analysis used both BEL-Pao (with and without glycoprotein) and representative BEL-Pao retroelements from the five branches of the BEL-Pao superfamily included in the Gypsy Database 2.0 (http://gydb.org).

The amino acid sequences for both analyses were aligned with the MAFFT algorithm (KATOH *et al.*, 2001) and automatically edited using Gblocks (CASTRESANA, 1999) with relaxed parameters (**Supplementary Material 2**). The most likely amino acid substitution model was determined using SMS (LEFORT; LONGUEVILLE; GASCUEL, 2017) on the ATCG online platform (http://www.atgc-montpellier.fr/).

Reconstruction of phylogenetic trees was carried out using MrBayes v3.2.2 x64 (RONQUIST; HUELSENBECK; TESLENKO, 2010), starting with three seeds and 1,000,000 generations. The convergence of the independent runs was detected when the standard deviation of all three seeds was lower than 0.05. The burn-in removed the first 25% of trees sampled and the remaining 75% were used to generate a posterior probability consensus tree and the phylogenies were visualized using iTOL (LETUNIC; BORK, 2007).

### PCR validation of Chuvirus solo-glycoproteins

Five solo glycoproteins without BEL-Pao signature were selected for PCR validation in three species available at the insectary of the Departamento de Entomologia - Instituto Aggeu Magalhães: *Aedes aegypti, Aedes albopictus* and *Culex quinquefasciatus*, as well as wild mosquitoes of the same species collected from different points at the Recife city (**Table 2**). The wild *Cx. quinquefasciatus* samples (Cxqui1301 and Cxqui1304) were collected in the Hospital das Clínicas - UFPE (8°02’51.9”S 34°56’45.6”W) during the years 2016 and 2017, as well as the wild sample Ae1471 of *Ae. aegypti*. The wild samples AlbZoo and AlbJB were collected in Parque Estadual Dois Irmãos (8°00’43.3”S 34°56’40.7”W) and Jardim Botânico do Recife (8°04’36.9”S 34°57’34.1”W) respectively, in 2017.

**Table 2:** Panorama of solo-glycoproteins signatures in different mosquito lineages.

|  | *Ae. albopictus* |  |  |  |  |  |  |  | *Ae. aegypti* |  |  |  | *Cx. quinquefasciatus* |  |  |  |  |  |
| --- | --- | --- | --- | --- | --- | --- | --- | --- | --- | --- | --- | --- | --- | --- | --- | --- | --- | --- |
|  | AalIns |  | AalC636 |  | AalZoo |  | AalJB |  | AaeIns |  | Aae 1471 |  | CxquiIns |  | Cxqui1304 |  | Cxqui1331 |  |
|  | L | R | L | R | L | R | L | R | L | R | L | R | L | R | L | R | L | R |
| AalFosh_15 | + | - | + | - | + | - | + | - |  |  |  |  |  |  |  |  |  |  |
| AalFosh_19 | + | - | - | - | + | - | + | - |  |  |  |  |  |  |  |  |  |  |
| AAeliv_06_L |  |  |  |  |  |  |  |  | - | + | - | + |  |  |  |  |  |  |
| AegAag2_12 |  |  |  |  |  |  |  |  | + | + | + | + |  |  |  |  |  |  |
| CquJoh_01 |  |  |  |  |  |  |  |  |  |  |  |  | - | + | - | + | - | + |

DNA extraction was performed from pools of 5 individuals (females and males) of each species with the protocol established by Ayres, 2003 (AYRES *et al.*, 2003), the quality and concentration of DNA was evaluated with Nanodrop 2000 (Thermo Fisher Scientific). Primers to amplify the endogenous gene of sodium channel (sodium channel protein para-like LOC109432678), were used as control to evaluate the DNA samples integrity before the EVE screening, and the regions that covers part of the EVE and a region of the mosquito genome were designed with Primmer3 (KORESSAAR; REMM, 2007) and validated with PrimmerBLAST (YE *et al.*, 2011) and OligoAnalyzer Tool (OWCZARZY *et* al., 2008) against the mosquitoes reference genomes (**Figure 1 A**). Primers information is shown in **Supplementary Material 7.** PCR was performed with GoTaq-Flexi G2 DNA-Polymerase following the Promega manufacturer’s protocol. All PCR reactions are conducted in a final volume of 25 uL containing 1 uL of each primer (10 uM), 2uL of dNTP (0.2 mM each base), 5.0 uL of GoTaq Flexi Buffer, 4.0 uL of MgCl2 (25 mM), 0.25 uL of GoTaq-Flexi G2 DNA-Polymerase, 3.0 uL of DNA sample and 8.75 uL of Ultrapure H20. The amplifications are conducted with the following program: Initial denaturation at 94°C for 2 minutes, 45 cycles at 94°C for 1 minute (denaturation), primer annealing at 51-59 °C for 50 seconds (depending on specific TM of each primer pair), extension at 72°C for 1 min followed by a final extension at 72 °C for 5 minutes. DNA amplification was visualized with 2% agarose gel stained with ethidium bromide and expected bands were extracted, purified and sequenced with DNA ABI Prism 3100 Genetic Analyser (Applied Biosystems) from both forward and reverse strands.

**Figure 1:**
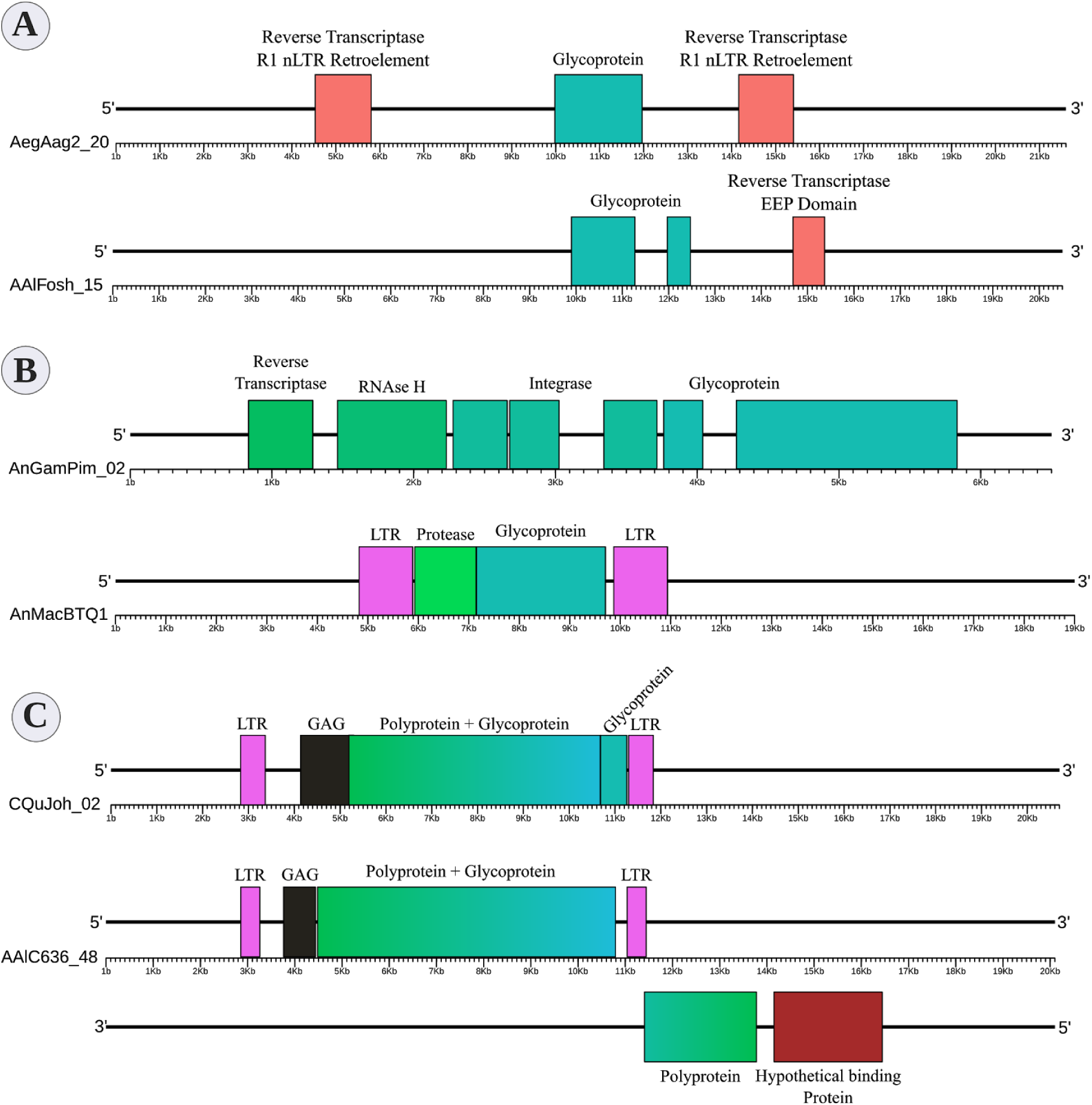
Example of glycoproteins of EVEs from *Chuviridae* family. **A**: Without the presence of BEL-Pao Retroelements; **B:** With the presence of Non-autonomous BEL-Pao Retroelements and **C:** Making part of Autonomous BEL-Pao Retroelements.

Forward and reverse electropherograms are analyzed with Geneious Prime version 2019.1.3 (KEARSE *et al.*, 2012) and consensus sequences were generated. Contigs from sequenced products are then aligned with EVEs identified by *in silico* analyses with Aliview (LARSSON, 2014).

## Results

### Supporting evidence of widespread chuvirus endogenization in different mosquito genomes

Four hundred and twenty-eight EVEs were identified in 32 out of 37 genomes screened. These elements corresponded to 409 glycoprotein fragments, 18 RNA-dependent RNA-polymerases (RdRp) and one nucleoprotein (**Supplementary Material 8**). After the exclusion of EVEs from small contigs and those in close proximity to uncertain sequences/assembled bases (NNNs), 279 sequences remained. Two hundred forty-one of these presented glycoprotein conserved amino acid domains.

In view of the previously described and confirmed abundance of chuvirus-derived glycoprotein EVEs, a more in-depth characterization of these EVEs was performed. Endogenous glycoproteins varied in size from 117 to 1977 nucleotides (median = 880) and amino acid length from 39 to 659 residues. The amino acid identity with each chuvirus genome used as a query varied from 29.03 to 56.41 percent (average identity = 32.83, SD = 6.31, **Supplementary Material 8**).

### Glycoproteins are mostly associated with elements from the BEL-Pao retrotransposon superfamily

Screening for BEL-Pao protein domain signatures and LTRs we found glycoproteins in three different contexts (**Figure 1, Table 1**). Thirty-seven glycoproteins were categorized as solo glycoproteins, since no BEL-Pao protein domain or LTR signature was found (**Figure 1 A**). Two hundred glycoproteins were flanked by BEL-Pao protein domains but with only one LTR or none, characterizing them as defective BEL-Pao elements (**Figure 1 B**). Sixty-three glycoproteins were associated with BEL-Pao domains flanked by two LTRs, characterizing them as potentially active elements. These included thirteen of those copies showing all domains and LTRs of complete BEL-Pao elements (**Figure 1 C**).

We also identified solo LTRs in seven genomes (**Table 1**). These varied in number from only one in the *Anopheles gambiae* Pimpirena assembly to 82 solo LTRs in the *Aedes albopictus* C636 assembly. Interestingly, the number of solo LTRs was found to be greater than the number of LTRs associated with full BEL-Pao elements in the *Aedes albopictus* C636 and *Anopheles maculatus* BTQ1 assemblies (**Table 1**).

### Endogenous and exogenous glycoproteins have similar 3D structures

Although BLAST and phylogenetic analysis indicate that the EVEs found share a common ancestor with chuviruses, the amino acid identity is considerably low (between 25 and 50%). Another way to obtain further support for the viral origin of these endogenous sequences is through 3D structure modeling. If these polypeptides resemble viral glyco/envelope proteins in 3D space, this would corroborate their viral origin.

The templates identified by the Phyre2 tool represent many type B glycoproteins homotrimmers from different viruses (**Figure 2**). It was possible to reconstruct three-dimensional models for all glycoproteins analyzed. All pdb files of modeled glycoproteins are available on **Supplementary Material 9** and Ramachandran Plots region values can be seen in **Supplementary Material 10**. Only two 3D models (AnMacBtQ1_01 and Wuhan Mosquito Virus 8) had more than 1.0% residues in disallowed regions. For all glycoproteins modeled the residues in core regions varied between 82.3% and 88.3% and in allowed regions from 9.3 to 14.2%.

**Figure 2:**
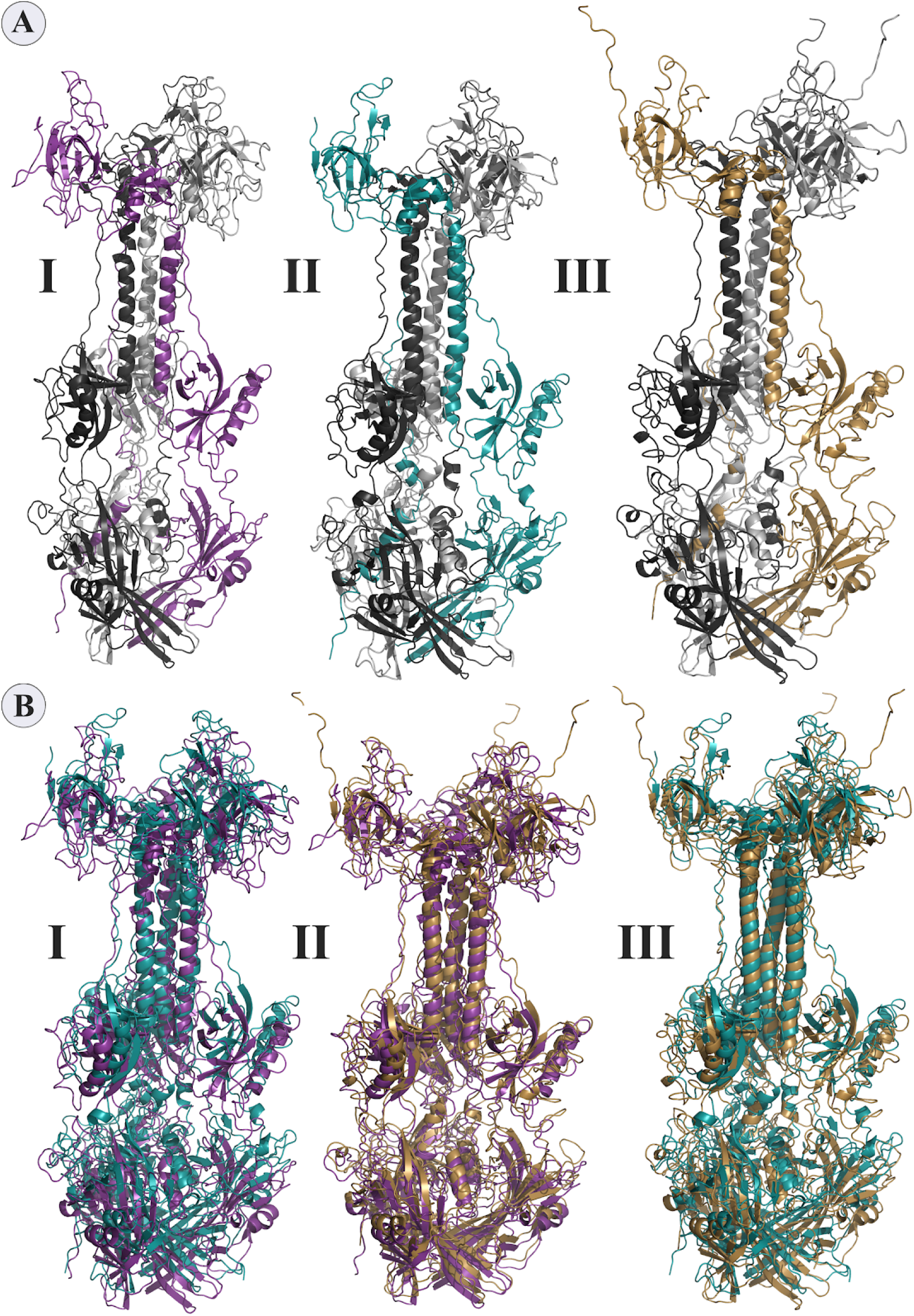
Molecular modeling of glycoproteins. **A:** Tridimensional models where **I** AegAag2_20 element, an example of solo-glycoprotein; **II** AAlC636_23 element, an example of glycoprotein fusioned with a complete Pao element; and **III** represents the glycoprotein of Mos8Chu0**. B:** Represents the tridimensional glycoproteins B alignments between: **I:** AegAag2_20 in purple and AAlC636_23 in blue, with TM-score equals to 0.79415; **II:** AegAag2_20 in purple and Mos8Chu0 in yellow, with TM-score equals to 0.76639; and **III:** AAlC636_23 in blue and Mos8Chu0 in yellow, with TM-score equals to 0.80812.

The TM-alignment between representatives of solo glycoproteins, glycoproteins fused with BEL-Pao elements and *bona fide* chuvirus glycoproteins demonstrates a similarity of three-dimensional structures greater than 0.7 (**Figure 2 B**), providing strong evidence that these glycoproteins are folded in similar way.

### Evidence of a Chuvirus ENV-like protein captured by elements from the BEL-Pao superfamily

*Bona fide* chuviruses with non segmented genomes (sequences in yellow) form a basal clade in the glycoprotein evolutionary tree (**Figure 3**), confirming the common origin of EVE glycoproteins and non segmented chuviruses. It is also worth noting the existence of a basal clade of *Aedes aegypti* EVEs clustered with one *bona fide* chuvirus - Mos8Chu0 (**Figure 4 A**).

**Figure 3:**
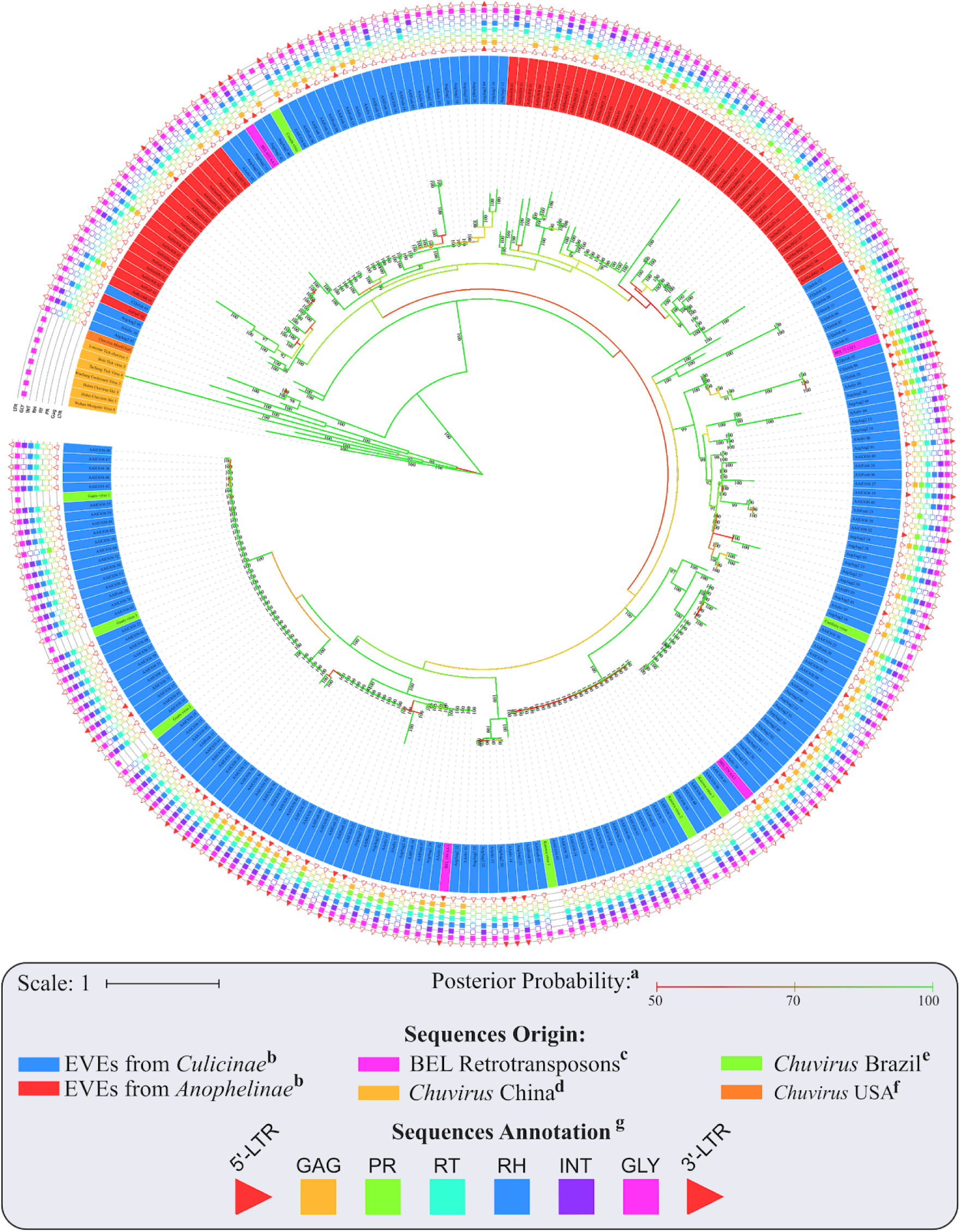
Phylogenetic representation of glycoproteins from *Chuviridae* family. Bayesian tree constructed after 5.000.000 generations from 3 seeds with standard deviation mean between final trees equal to 0.02. **a:** Posterior probability in percentage, only values greater than 90 was plotted on phylogeny; **b:** Sequences identified as EVEs derived from *Chuviridae* family; **c:** Retrotransposons of BEL-Pao superfamily available on RepBase and that showed similarity with chuviruses glycoproteins; **d:** Chuviruses identified in China by Shi et al 2016; **e:** Sequences described in Brazil as chuviruses by Pinto, A. et al 2017; **f:** Chuvirus available on NCBI (access KX924631.1); **g:** LTR = Long Terminal Repeat, GAG = Capsid protein; PR = Protease, RT = Reverse Transcriptase, RH = RNAse H, Integrase and GLY = Glycoprotein. Phylogeny available on: https://itol.embl.de/tree/200133261329821563538693.

**Figure 4:**
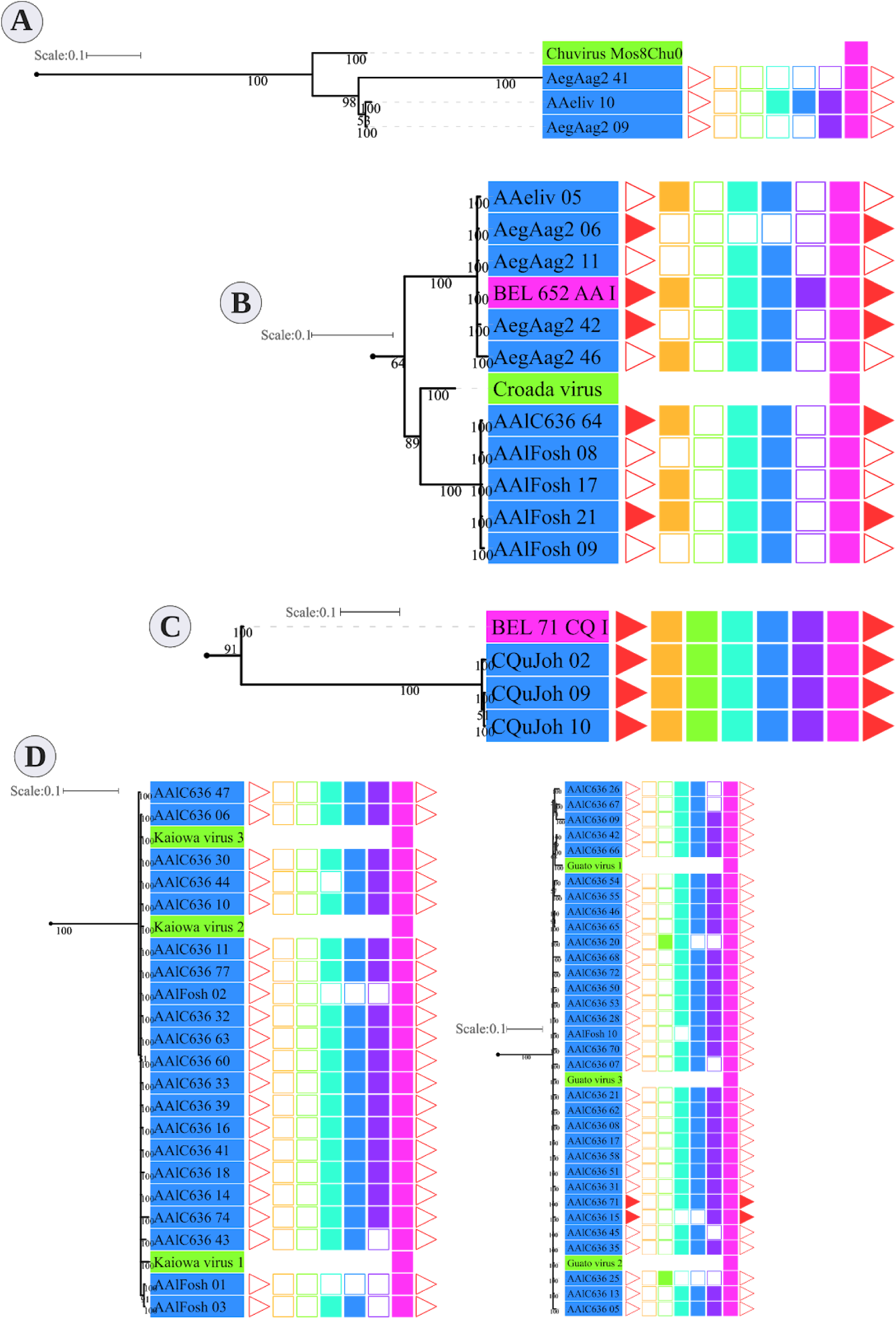
Zoom of 4 clades from Figure 4. **A:** Basal clade with *bona fide* chuvirus and EVEs; **B:** Clades with copies of the same elements; **C:** Clade with EVEs and BEL-Pao Retrotroelement; **D:** Clades with EVEs and chuvirus from Pinto, A. et al 2017.

Three main findings should be reported regarding EVE clades: i) the presence of various high-identical EVE copies, indicated by clades containing several sequences with near-zero branch lengths (**Figure 4 B**); ii) the presence of glycoproteins inside BEL-Pao retroelements from RepBase intervening into various EVE clades (**Figure 4 C**); and iii) the presence of sequences described as *bona fide* chuviruses (LARA PINTO *et al.*, 2017; PAUVOLID-CORRÊA *et al.*, 2016) inside and outside of clades mostly composed of EVE copies (**Figure 4 D**).

The evolutionary tree using the RT and RH regions from both BEL-Pao retroelements, including elements containing or not chuvirus glycoproteins, shows four clearly defined clades (**Figure 5**). One of these has 181 elements without glycoproteins closely-related to BEL elements in Branch 1 (**Figure 5**). The other three are closely-related clades with Pao elements from Branch 2, of which 129 are composed of elements with glycoproteins (red scalene triangles) and 122 of elements without glycoproteins (blue scalene triangles, **Figure 5**). It is clear that chuviruses glycoproteins are specifically associated with elements from the Pao family - the now called Anakin elements.

**Figure 5:**
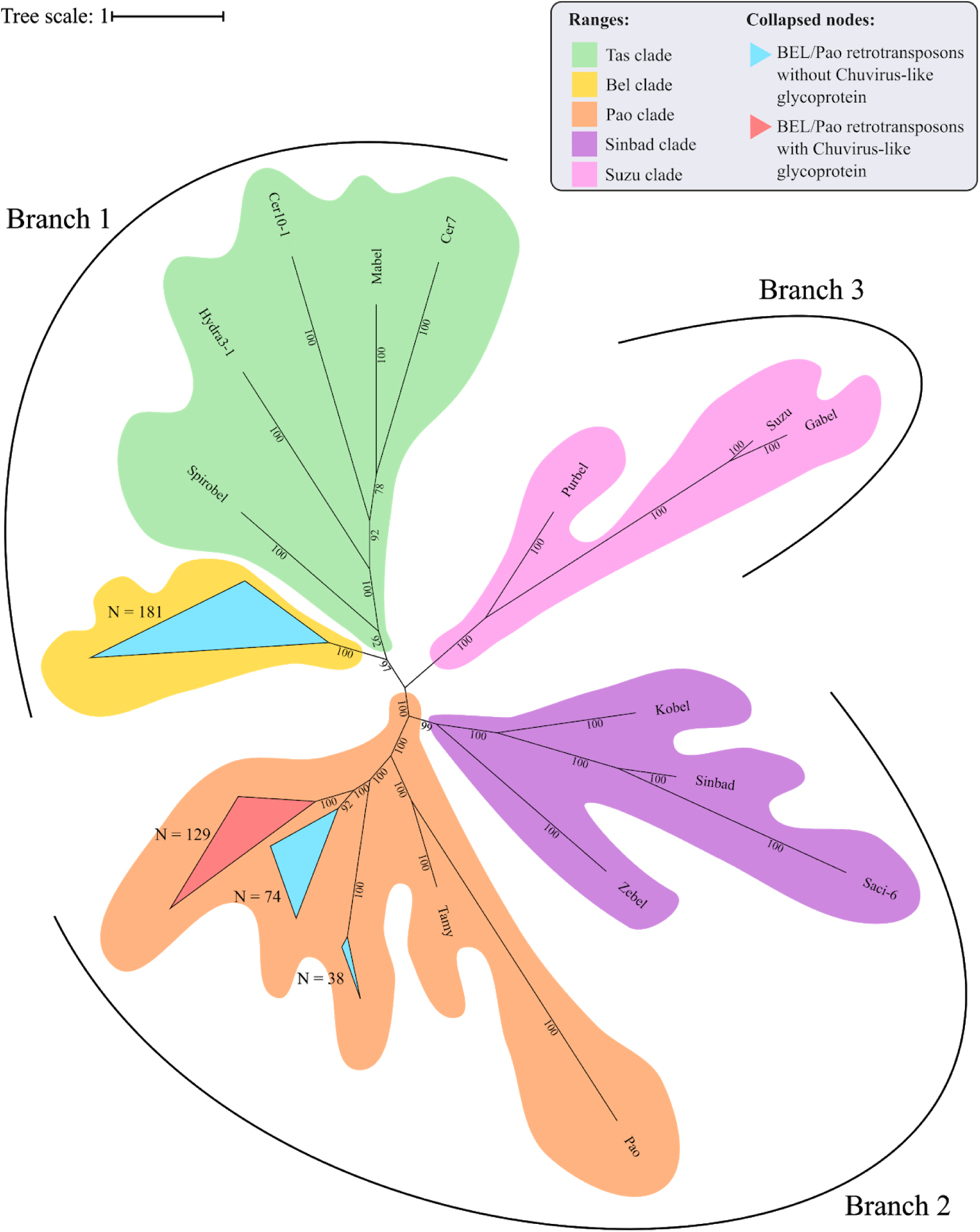
Phylogenetic representation of Reverse Transcriptase and RNAse H regions of Polyproteins from BEL-Pao Retroelements. Bayesian tree constructed after 1.000.000 generations from 3 seeds with standard deviation mean between final trees equal to 0.03. Support values in posterior probability. Branches and clades are annotated according BEL/Pao Retroelements phylogeny patterns available on: http://gydb.org/index.php/. Uncollapsed phylogeny with domain annotation is present on https://itol.embl.de/tree/200133261405581567779232.

There is a large number of Pao elements without glycoproteins, but comparing it with the Anakin elements the last are more abundant and widespread, in a larger number of mosquito species, than Pao elements without glycoproteins. (**Supplementary Material 11**).

### PCR validation of solo-glycoproteins in different mosquito lineages

Five primer pairs were designed to amplify both chuvirus/mosquito integration junction of solo-glycoprotein regions. Four lineages of *Ae. albopictus*, two lineages of *C. quinquefasciatus* and two lineages of *Ae. aegypti* were screened (**Table 2, Supplementary Material 12**). At least one EVE-mosquito boundaries were sequenced successfully for all five solo glycoproteins (**Table 2**), showing a consistent alignment to the corresponding genomic region in the available genomes (**Supplementary Material 12**). This confirms the integration of the EVEs and highlights that these insertions were conserved and an ancient component of these genomes.

## Discussion

Viruses and transposable elements share one or few common ancestors and hence present similar features, particularly for some retroviruses and retrotransposons that are differentiated by the presence/absence of an envelope gene (glycoprotein). The envelope protein is responsible for the infection capacity of the former (HAYWARD, 2017). But more than that, these entities exchange genetic information between them and with their host genome (DREZEN *et al.*, 2017; FRANK; FESCHOTTE, 2017; GILBERT; CORDAUX, 2017; SINHA; JOHNSON, 2017). Several instances are of interexchange of functional genes that allowed major changes in their evolutionary history as known such as the acquisition of a complete herpesvirus by a piggyBac transposon (INOUE *et al.*, 2017) or the emergence of retroviruses by envelope capture (KIM *et al.*, 2004). Here, we characterized a new event of gene sequence exchange between viruses, the mosquito host genome and retrotransposons with the acquisition of a glycoprotein (envelope) gene from a chuvirus by a Pao retrotransposon (named Anakin). Besides, we also found evidence of past and likely current arms race between these entities.

The *Chuviridae* family was first described in 2016 by Shi et al. in several arthropod species (including mosquitoes) through metatranscriptomic sequencing (SHI *et al.*, 2016). chuviruses have three possible genomic structures: a complete or bi-segmented circular genome, identified in ticks, crab, flies, spiders, cockroaches and mosquitoes, and a linear structure, identified in flies and crabs. In the bi-segmented structure, the glycoprotein gene is always flanked downstream by a nucleoprotein and by a viral particle protein (SHI *et al.*, 2016). In the same study, the authors identified endogenous chuvirus elements in both mosquito and other insect genomes. Two subsequent studies also identified EVEs derived from chuvirus in mosquito genomes (RUSSO *et al.*, 2019; WHITFIELD *et al.*, 2017) but focused only on the *Aedes aegypti* genome. Whitfield et al. 2017 characterized two interesting features of chuvirus-derived EVEs from *A. aegypti*: i) large differences among endogenized genomic regions, with several glycoproteins and few RdRps and nucleoproteins; and ii) enrichment of BEL-Pao retroelements around these EVEs (RUSSO *et al.*, 2019; WHITFIELD *et al.*, 2017).

Virus genome replication is tightly linked to virus genome structure. Replication origin and orientation may favor the endogenization of the genomic regions that are first copied/transcribed, since these regions are produced more abundantly than the last replicated/transcribed regions (RUSSO *et al.*, 2019; WHITFIELD *et al.*, 2017). Although little is known about chuvirus replication, the position of the genes in non-segmented genomes suggests that endogenization should occur more frequently for nucleoproteins or RdRp (in the terminal regions of the virus genomes) than for glycoproteins (around the middle of the virus genome). However, for segmented genomes, each segment could be integrated independently and hence one should expect to find a similar amount of different integrated viral genomic regions. We detected more glycoproteins than nucleoproteins and RdRps regions integrated into the mosquito genomes. This endogenization pattern is not expected to any segmented or non-segmented Chuvirus genomes and hence does not allow us to reach a conclusion about the genomic structure of the original chuvirus genome. But, some additional evidence points to another more likely explanation to explain the glycoprotein discrepancy. We detected that many of these glycoproteins are integrated into potentially active BEL-Pao retrotransposons, more specifically into retroelements from the Pao family, suggesting that the high glycoprotein copy number is a result of the replication of Pao retrotransposons and that this protein may be an integral part of these elements as an envelope gene.

Acquisition and loss of envelope proteins have been detected in a number of viruses and retrotransposons, blurring the distinction between these entities. Substantial evidence already exists showing that many retroviruses can turn into an intragenomic lifestyle when the ENV protein is lost or becomes defective as a result of mutations and that various retroviruses have originated from retrotransposons (intragenomic lifestyle) that acquired new envelope genes (MALIK, H. S.; HENIKOFF; EICKBUSH, 2000). Specifically for BEL-Pao retrotransposons, there are three well studied examples of ENV-like protein acquisition by active retroelements (MALIK, H. S.; HENIKOFF; EICKBUSH, 2000). The *Tas* retrotransposon from *Ascaris lumbricoides* acquired an ENV protein from *Phlebovirus* (FELDER *et al.*, 1994). *The Cer* retrotransposon from *Caenorhabditis elegans* acquired an ENV protein from Herpesvirus (BROWNING *et al.*, 1996) and an ENV was acquired from a Gypsy retrovirus by a Roo retrotransposon (MALIK, HARMIT S.; HENIKOFF, 2005). We describe here a fourth event, involving Pao retrotransposons and glycoproteins of chuvirus, supported by the strong phylogenetic association between glycoproteins of Pao retrotransposons and EVEs from the *Chuviridae* family and the identification of glycoproteins inside complete Pao structures flanked by LTRs. This recombination event associated with further retrotransposition of Anakin explains the high abundance of glycoproteins inside mosquito genomes when compared with the other chuvirus genes.

The distribution of EVEs derived from chuvirus in mosquito species from both the Culicinae and Anophelinae subfamilies dates the integration of chuvirus glycoproteins into the ancestor of the Culicidae family, around 190 million years ago (HEDGES; DUDLEY; KUMAR, 2006). The endogenization of a chuvirus glycoprotein may have occurred directly into a Pao retrotransposon, thereby giving rise to the Pao retrovirus (**Figure 6A**). Alternatively, this hybrid element may have emerged from a recombination event involving a Pao retroelement and a chuvirus-derived solo glycoprotein after its endogenization (**Figure 6A**). After viral envelope protein endogenization into the host genome, two major events may occur: exaptation or molecular domestication of the viral protein leading to a new host function or the emergence of an antiviral mechanism. The domesticated viral envelope may be selected as a countermeasure against cognate circulating virus to effectively prevent new virus particles from entering the cell. For instance, host endogenous envelope proteins may be produced and exported to the extracellular space by the host cells. These polypeptides will then compete for the host cell receptors with circulating viruses (JOHNSON, 2019; ARMEZZANI *et al.*, 2014; ITO *et al.*, 2013; ROBINSON *et al.*, 1981). The similarity of solo glycoprotein and glycoproteins from *bona fide* chuviruses and the Anakin retrovirus, as shown by sequence identity (**Supplementary Material 13**) and comparison of three-dimensional structures (**Figure 4A**), indicates that some of these endogenous glycoproteins may play a role in antiviral defense mechanisms against circulating chuviruses or even against Anakin retroviruses through competition for cell receptors (**Figure 6 B**). This hypothesis is corroborated by the conservation of several solo glycoprotein integration in different populations of *Ae. aegypti, Ae. albopictus* and *Cx. quinquefasciatus* species (**Table 2**). But we can not rule out the repurposing of more divergent glycoproteins to other unknown functions at the host level.

**Figure 6:**
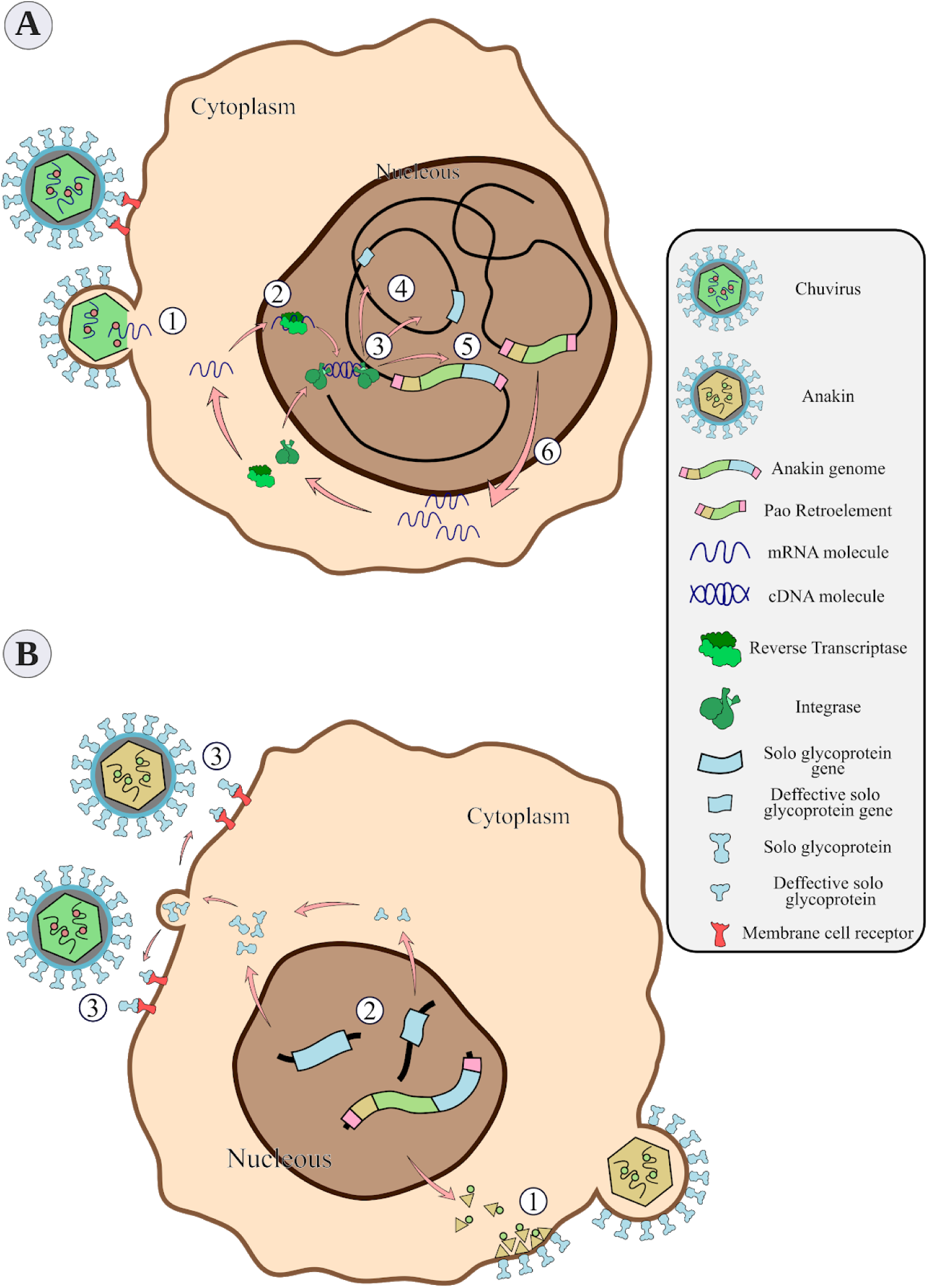
General scheme of the chuvirus derived EVE integration into the germinative cells of the host genome, phusion with *Pao* retroelements and potential antiviral mechanisms by receptor competition. A - Bonafide circulating chuvirus infecting a host cell and realising their single strand RNA into the cell cytoplasm (1); RT from retrotransposons fortuitously recognize the viral genome and retro transcribe to dsDNA (2); dsDNA is directly integrated into the host genome by double strand break repair mechanisms or by recognition and integration using integrase proteins from retroelements (3) either in different genomic loci (4) or into a *Pao* retroelement boundaries (5); Expression of Pao proteins (6); B - chuvirus-derived glycoproteins are dispersed in many genomic loci but have been replicated substantially by the *Pao* + chuvirus glycoprotein element (Anakin). Anakin retrovirus can assemble viral particles with envelope proteins allowing it to infect new host cells (1); Complete or defective solo glycoprotein are translated and exported to the extracellular environment (2) and can block new viral infection of both Anakin and chuvirus by binding to cell membrane virus protein receptors (3).

Finally, our results show the importance of taking EVEs into account in metaviromic studies (NOURI *et al.*, 2018). Some of the chuvirus EVEs detected in our study exhibited a high degree of identity (99.08 to 100%) with previously described circulating chuviruses *Kaiowa, Guato, Cumbaru* and *Croada* (LARA PINTO et al., 2017; PAUVOLID-CORRÊA et al., 2016). Although, we cannot rule out the possibility that some of these sequences may indeed have come from circulating chuviruses that infect *Ae. albopictus* species, the high identity values of these “viruses” with the EVEs found here strongly suggest that these previously defined chuviruses are in fact endogenous elements of the *Ae. albopictus* genome (**Supplementary Material 14**).

## Conclusion

In this study we revealed the diversity of endogenous virus elements derived from the *Chuviridae* family in mosquito genomes. Our results show that such EVEs are widely distributed across the *Culicidae* family and are possibly involved in two major processes: the replication of Pao retroelements, through acquisition of chuvirus glycoproteins; and a possible antiviral response against *bona fide* chuviruses and Anakin retroviruses originating from a fusion of Pao retroelements and the chuvirus envelope gene. These results shed new light on the dynamic evolution of EVEs and retrotransposons inside the mosquito genomes and point to the need for further studies regarding the role of these elements at the protein level against *bona fide* viruses, retroviruses and at the mosquito biology.

## Declarations

## List of abbreviations

EVEs: Endogenous Virus Elements
LTR: Long Terminal Repeats
piRNA: Piwi-interacting RNA
RdRp: RNA-dependent RNA polymerase
NCBI: National Center for Biotechnology Information
kb: kilobase
kbp: kilobase pair
ORF: Open Reading Frame

## Ethics approval and consent to participate

Not applicable.

## Consent for publication

Not applicable.

## Authors’ contributions

G.L.W. conceive the study. G.L.W and A.M.R. planned and supervised the work. F.Z.D. carried out the genomics analysis and wrote the manuscript. C.S.V. carried out the molecular modeling analysis. All authors agreed with the final version of the manuscript.

## Acknowledgments

We thank the bioinformatics core at the Instituto Aggeu Magalhães (IAM) for technical assistance. We thank the collaborators from the Departamento de Entomologia - Instituto Aggeu Magalhães for the wild mosquitoes samples.

## Supplementary Material

All supplementary material was available on: 10.6084/m9.figshare.11336258

## REFERENCES

Altschul, Stephen F. et al. Basic local alignment search tool. Journal of molecular biology v. 215, n. 3, p. 403–10, 1989.

Armezzani, Alessia et al. “Ménage à Trois”: the evolutionary interplay between JSRV, enJSRVs and domestic sheep. Viruses v. 6, n. 12, p. 4926–4945, 9 dez. 2014.

Ayres, C. F. J. et al. Genetic differentiation of Aedes aegypti (Diptera: Culicidae), the major dengue vector in Brazil. Journal of medical entomology v. 40, n. 4, p. 430–435, 1 jul. 2003.

Browning, H. et al. Macrorestriction analysis of Caenorhabditis elegans genomic DNA. Genetics v. 144, n. 2, p. 609–619, 1 out. 1996.

Castresana, J. Selection of conserved blocks from multiple alignments for their use in phylogenetic analysis. Molecular biology and evolution v. 17, n. 4, p. 540–552, 1999.

Drezen, Jean-Michel et al. Endogenous viruses of parasitic wasps: variations on a common theme. Current opinion in virology v. 25, p. 41-48, 17 ago. 2017.

Felder, H. et al. Tas, a retrotransposon from the parasitic nematode Ascaris lumbricoides. Gene v. 149, n. 2, p. 219–225, 18 nov. 1994.

Feschotte, Cédric; Gilbert, Clément. Endogenous viruses: insights into viral evolution and impact on host biology. Nature Reviews Genetics v. 13, p. 283–296, 2010.

Forterre, Patrick; Prangishvili, David. The great billion-year war between ribosome- and capsid-encoding organisms (cells and viruses) as the major source of evolutionary novelties. Annals of the New York Academy of Sciences v. 1178, p. 65–77, 1 out. 2009.

Frank, John A.; Feschotte, Cédric. Co-option of endogenous viral sequences for host cell function. Current opinion in virology v. 25, p. 81-89, 16 ago. 2017.

Gel, Bernat; Serra, Eduard. karyoploteR: an R/Bioconductor package to plot customizable genomes displaying arbitrary data. Bioinformatics (Oxford, England) v. 33, n. 19, p. 3088–3090, 1 out. 2017.

Gilbert, Clément; Cordaux, Richard. Viruses as vectors of horizontal transfer of genetic material in eukaryotes. Current opinion in virology v. 25, p. 16-22, 30 ago. 2017.

Goubert, Clément et al. De novo assembly and annotation of the Asian tiger mosquito (Aedes albopictus) repeatome with dnaPipeTE from raw genomic reads and comparative analysis with the yellow fever mosquito (Aedes aegypti). Genome biology and evolution v. 7, n. 4, p. 1192–1205, 11 mar. 2015.

Hayward, Alexander. Origin of the retroviruses: when, where, and how? Current opinion in virology v. 25, p. 23-27, 30 ago. 2017.

Hedges, S. Blair; DUDLEY, Joel; KUMAR, Sudhir. TimeTree: a public knowledge-base of divergence times among organisms. Bioinformatics (Oxford, England) v. 22, n. 23, p. 2971–2972, 1 ez. 2006.

Inoue, Yusuke et al. Complete fusion of a transposon and herpesvirus created the Teratorn mobile element in medaka fish. Nature communications v. 8, n. 1, p. 551, 15 set. 2017.

ITO, Jumpei et al. Refrex-1, a soluble restriction factor against feline endogenous and exogenous retroviruses. Journal of virology v. 87, n. 22, p. 12029–12040, 21 nov. 2013.

Johnson, Welkin E. Origins and evolutionary consequences of ancient endogenous retroviruses. Nature reviews. Microbiology v. 17, n. 6, p. 355–370, 1 jun. 2019.

Katoh, K. et al. MAFFT: a novel method for rapid multiple sequence alignment based on fast Fourier transform. Nucleic Acids Research v. 30, n. 14, p. 3059–3066, 2001.

Katzourakis, Aris; GIFFORD, Robert J. Endogenous viral elements in animal genomes. PLoS genetics v. 6, n. 11, p. e1001191, 18 nov. 2010.

Kearse, Matthew et al. Geneious Basic: an integrated and extendable desktop software platform for the organization and analysis of sequence data. Bioinformatics (Oxford, England) v. 28, n. 12, p. 1647–1649, 15 jun. 2012.

Kelley, Lawrence A. et al. The Phyre2 web portal for protein modeling, prediction and analysis. Nature protocols v. 10, n. 6, p. 845–858, 7 jun. 2015.

Kim, Felix J. et al. Emergence of vertebrate retroviruses and envelope capture. Virology v. 318, n. 1, p. 183–191, 5 jan. 2004.

Koressaar, T.; Remm, M. Enhancements and modifications of primer design program Primer3. Bioinformatics v. 23, n. 10, 15 maio 2007.

LARA Pinto, Andressa Zelenski De et al. Novel viruses in salivary glands of mosquitoes from sylvatic Cerrado, Midwestern Brazil. PloS one v. 12, n. 11, p. e0187429, 8 nov. 2017.

Larsson, A. AliView: a fast and lightweight alignment viewer and editor for large datasets. Bioinformatics v. 30, n. 22, 15 nov. 2014.

Lefort, Vincent; Longueville, Jean-Emmanuel; Gascuel, Olivier. SMS: Smart Model Selection in PhyML. Molecular biology and evolution v. 34, n. 9, p. 2422–2424, 1 set. 2017.

Letunic, Ivica; BORK, Peer. Interactive Tree Of Life (iTOL): an online tool for phylogenetic tree display and annotation. Bioinformatics (Oxford, England) v. 23, n. 1, p. 127–128, 1 jan. 2007.

LI, Weizhong; GODZIK, Adam. Cd-hit: a fast program for clustering and comparing large sets of protein or nucleotide sequences. Bioinformatics v. 22, p. 1658-1659, 2005.

Llorens, Carlos et al. The Gypsy Database (GyDB) of mobile genetic elements: release 2.0. Nucleic acids research v. 39, n. Database issue, p. D70-D74, 29 jan. 2011.

Malik, H. S.; Henikoff, S.; Eickbush, T. H. Poised for contagion: evolutionary origins of the infectious abilities of invertebrate retroviruses. Genome research v. 10, n. 9, p. 1307–1318, 1 set. 2000.

Malik, Harmit S.; Henikoff, Steven. Positive selection of Iris, a retroviral envelope-derived host gene in Drosophila melanogaster. PLoS genetics v. 1, n. 4, p. e44, 1 out. 2005.

Marchler-Bauer, A. et al. CDD: a Conserved Domain Database for the functional annotation of proteins. Nucleic Acids Research v. 39, n. Database, 1 jan. 2011.

Nene, V. et al. Genome sequence of Aedes aegypti, a major arbovirus vector. Science v. 316, n. 5832, p. 1718–1723, 2006.

Nouri, Shahideh et al. Insect-specific viruses: from discovery to potential translational applications. Current opinion in virology v. 33, p. 33-41, 23 ez. 2018.

Okonechnikov, K.; Golosova, O.; Fursov, M. Unipro UGENE: a unified bioinformatics toolkit. Bioinformatics v. 28, n. 8, 15 abr. 2012.

Owczarzy, Richard et al. IDT SciTools: a suite for analysis and design of nucleic acid oligomers. Nucleic acids research v. 36, n. Web Server issue, p. W163-W169, 1 jul. 2008.

Palatini, Umberto et al. Comparative genomics shows that viral integrations are abundant and express piRNAs in the arboviral vectors Aedes aegypti and Aedes albopictus. BMC genomics v. 18, n. 1, p. 512, 5 jul. 2017.

Pauvolid-Corrêa, Alex et al. Novel Viruses Isolated from Mosquitoes in Pantanal, Brazil. Genome announcements v. 4, n. 6, 3 nov. 2016.

Robinson, H. L. et al. Host Susceptibility to endogenous viruses: defective, glycoprotein-expressing proviruses interfere with infections. Journal of virology v. 40, n. 3, p. 745–751, 1 ez. 1981.

Ronquist, Fredrik; Huelsenbeck, John; Teslenko, Maxim. Draft Mr. Bayes version 3.2 Manual. [S.l: s.n.]., 2010

Russo, Alice G. et al. Novel insights into endogenous RNA viral elements in and other arbovirus vector genomes. Virus evolution v. 5, n. 1, p. vez010, 18 jan. 2019.

Shi, Mang et al. Redefining the invertebrate RNA virosphere. Nature v. 540, n. 7634, p. 539–543, 22 ez. 2016.

Sinha, Anindita; Johnson, Welkin E. Retroviruses of the RDR superinfection interference group: ancient origins and broad host distribution of a promiscuous Env gene. Current opinion in virology v. 25, p. 105-112, 19 ago. 2017.

Théron, Emmanuelle et al. Distinct features of the piRNA pathway in somatic and germ cells: from piRNA cluster transcription to piRNA processing and amplification. Mobile DNA v. 5, n. 1, 2013.

Weiss, Robin A. Exchange of Genetic Sequences Between Viruses and Hosts. Current topics in microbiology and immunology v. 407, p. 1–29, 2016.

Whitfield, Zachary J. et al. The Diversity, Structure, and Function of Heritable Adaptive Immunity Sequences in the Aedes aegypti Genome. Current biology: CB v. 27, n. 22, p. 3511-3519.e7, 20 nov. 2017.

XU, Zhao; Wang, Hao. LTR_FINDER: an efficient tool for the prediction of full-length LTR retrotransposons. Nucleic acids research v. 35, n. Web Server issue, p. W265-W268, 7 jul. 2007.

YE, Jian et al. Primer-BLAST: a tool to design target-specific primers for polymerase chain reaction. BMC bioinformatics v. 13, p. 134, 2011.

